# Flexible pri-miRNA structures enable tunable production of 5’ isomiRs

**DOI:** 10.1101/2021.08.18.456839

**Authors:** Xavier Bofill-De Ros, Zhenyi Hong, Ben Birkenfeld, Sarangelica Alamo-Ortiz, Acong Yang, Lisheng Dai, Shuo Gu

## Abstract

Drosha cleavage of a pri-miRNA defines mature microRNA sequence. Drosha cleavage at alternative positions generates 5’ isoforms (isomiRs) which have distinctive functions. To understand how pri-miRNA structures influence Drosha cleavage, we performed a systematic analysis of the maturation of endogenous pri-miRNAs and their variants both *in vitro* and *in vivo*. We show that, in addition to previously known features, the overall structural flexibility of pri-miRNA impacts Drosha cleavage fidelity. Internal loops and nearby G·U wobble pairs on the pri-miRNA stem induce the use of non-canonical cleavage sites by Drosha, resulting in 5’ isomiR production. By analyzing patient data deposited in The Cancer Genome Atlas, we provide evidence that alternative Drosha cleavage of pri-miRNAs is a tunable process that responds to the level of pri-miRNA-associated RNA-binding proteins. Together, our findings reveal that Drosha cleavage fidelity can be modulated by altering pri-miRNA structure, a potential mechanism underlying 5’ isomiR biogenesis in tumors.

**HIGHLIGHTS:**

- Flexible pri-miRNA structures lead to 5’ isomiR production
- Internal loops and G·U pairs of pri-miRNA contribute to alternative Drosha cleavages
- Alternative Drosha cleavage results in 5’ isomiRs from both strands of pre-miRNAs
- 5’ isomiR production is upregulated by pri-miRNA-associated RBPs in cancers

**GRAPHICAL ABSTRACT:** 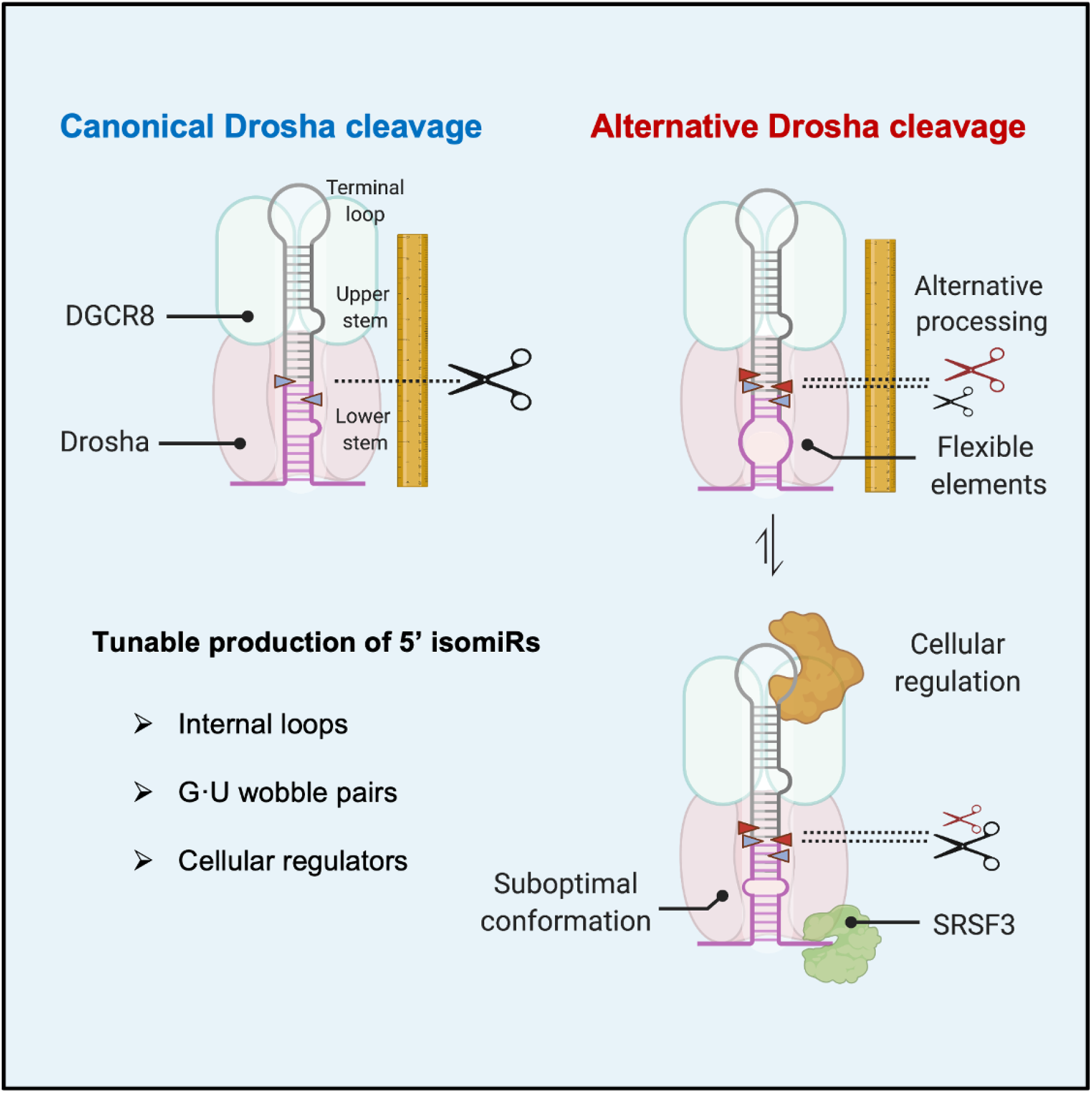

## INTRODUCTION

MicroRNAs (miRNAs) play critical roles in cellular physiology by inhibiting gene expression via post-transcriptional mechanisms (Gebert and MacRae, 2019). Misregulation of miRNAs is involved in the pathogenesis of many diseases including cancer (Lin and Gregory, 2015). MiRNAs are transcribed as part of longer transcripts, primary miRNA (pri-miRNAs). Pri-miRNAs fold into hairpin structures, which are recognized by RNase III enzymes Drosha and Dicer (Ha and Kim, 2014). After stepwise endonucleolytic cleavage, the resulting RNA duplexes (~22 nt) are loaded onto Argonaute proteins. Eventually, one of two strands remains associated with the Argonaute protein, forming the core of the RNA-induced silencing complex (RISC) (Khvorova et al., 2003; Schwarz et al., 2003). In metazoans, miRNA guides RISC to target mRNAs by partial base-pairing, providing inhibition specificity (Iwakawa and Tomari, 2015; Jonas and Izaurralde, 2015). The sequence of nucleotides from position 2 to 7, counting from the miRNA 5’ end, plays a crucial role in this process. Base-pairing of this small “seed” region with targets is required and often sufficient for a miRNA to function (Bartel, 2009). Ends of miRNAs are defined by Drosha and Dicer cleavages. Drosha cutting at an alternative position generates miRNA isoforms (isomiRs) with a distinct 5’ end and an altered seed sequence (Figure 1A). As these 5’ isomiRs have an altered target repertoire, alternative Drosha cleavage profoundly impacts miRNA function. This highlights the importance of understanding the mechanisms by which Drosha cleavage fidelity is governed.

**Figure 1.**
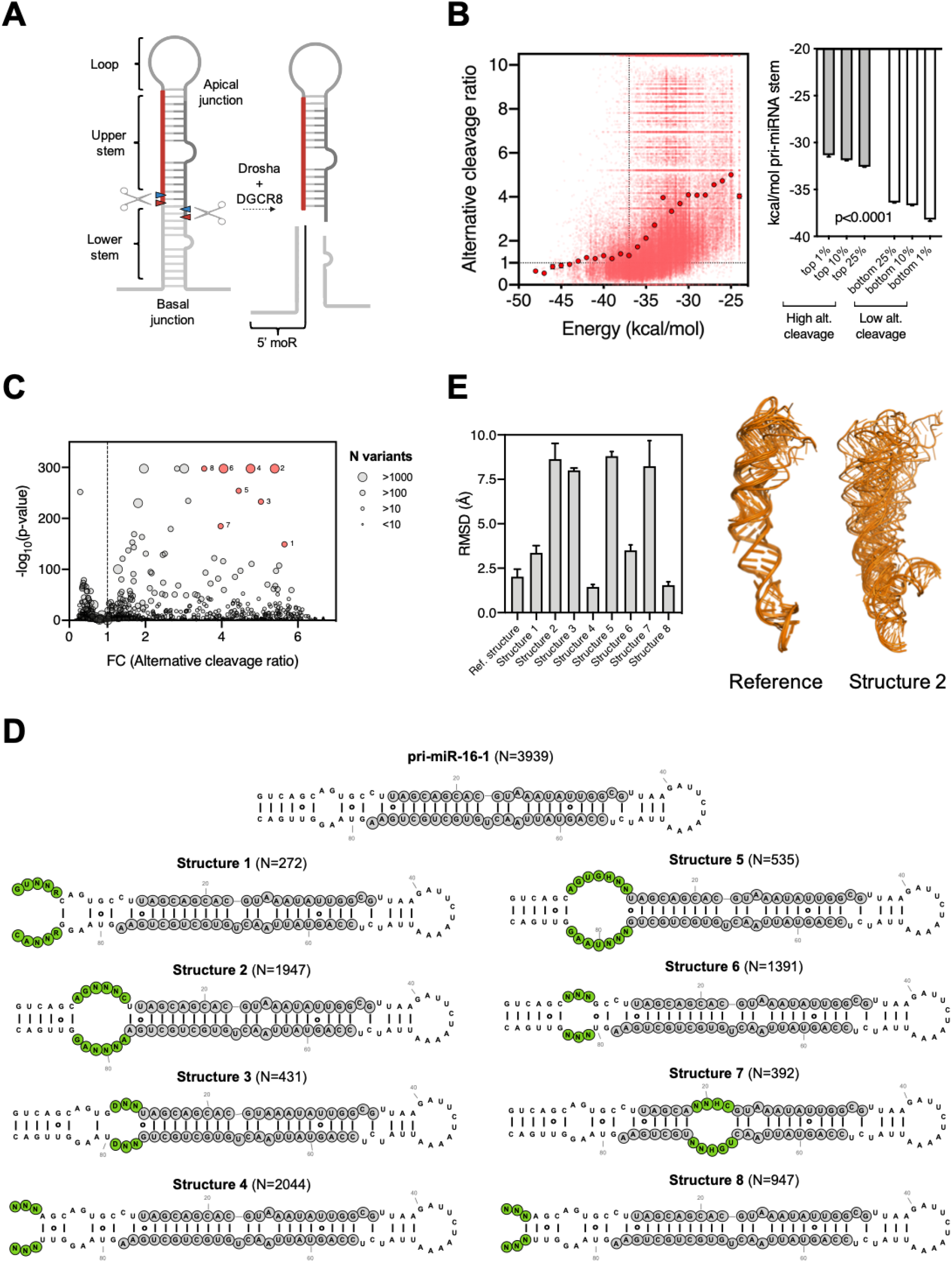
More flexible pri-miRNA structure correlates with alternative Drosha cleavages. (A) Scheme of the pri-miRNA structure. Red and blue arrows indicate Drosha cleavage sites which can be measured from the sequence of the 5’ moR products. (B) Left, scatter plot of the alternative cleavage ratio (relative to pri-miR-16-1 wild-type sequence) as a function of the minimum folding energy (kcal/mol) of each sequence variant. Right, minimum folding energy (kcal/mol) of pri-miR-16-1 sequence variants with high and low alternative cleavage levels. (C) Average alternative cleavage ratios of pri-miR-16-1 sequence variants with the same predicted secondary structure were plotted against the p-values calculated by comparison with the variants with the same secondary structure as the pri-miR-16-1 wild-type (N=3939) using the Wilcoxon test. The size of the dots indicates the number of sequence variants supporting that structure group. (D) Structural variants with the most significant increases in alternative Drosha cleavage compared to variants with the native structure. N indicates the number of sequence variants supporting that structure group. Nucleotides involved in the structural feature are indicated in green and shown in the IUPAC nucleotide code. (E) RMSD (Å) of 20 sequence variants from each structural group against pri-miR-16-1 wild-type. Tertiary structures were predicted using RNAComposer.

Drosha and its cofactor DGCR8 form a complex called Microprocessor, which determines its cleavage site by recognizing a set of structural and sequence features of pri-miRNA. The pri-miRNA hairpin is structurally defined by a terminal loop (8-38nt), a stem with a high degree of complementarity (~35bp) and unstructured flanking sequences (Fromm et al., 2015). The stem is divided into an upper stem where the mature miRNA sequence resides and a lower stem next to the flanking regions (Nguyen et al., 2015). Drosha senses the basal junction between the flanking regions and the lower stem, and cleaves ~11bp away (Han et al., 2006). DGCR8 binds the apical junction between the upper stem and terminal loop, facilitating the recognition of pri-miRNAs (Nguyen et al., 2015; Zeng et al., 2005). Although the Drosha-DGCR8 complex is sufficient to process most pri-miRNAs *in vitro (Rice et al., 2020)*, recent findings indicate that Drosha cleavage is modulated by a large number of pri-miRNA associated RNA-binding proteins (RBPs) *in vivo (Treiber et al., 2017)*. Several sequence motifs, including a “UG” at the basal junction, a “UGU” at apical junction and “CNNC” at the flanking region, are recognized by Drosha, DGCR8 and SRSF3, respectively (Auyeung et al., 2013; Fang and Bartel, 2015; Kwon et al., 2019).

While all these structural and sequence features contribute to efficient Drosha processing, the extent to which these features impact Drosha cleavage fidelity remains unclear. Several studies have shown that stem length affects Drosha cleavage fidelity by defining the relative distances between the expected cleavage site, the basal junction, and the apical junction of a pri-miRNA (Burke et al., 2014; Ma et al., 2013; Roden et al., 2017). Furthermore, an interaction between Drosha dsRBD and a mGHG motif in the lower stem promotes precise cleavage by facilitating the alignment between Drosha and pri-miRNA substrates (Fang and Bartel, 2015; Kwon et al., 2019). More recently, mismatched and wobble base pairs along the upper stem were found to impact Drosha cleavage fidelity as well (Li et al., 2020). However, these mechanisms individually or in combination cannot fully account for the extent of alternative Drosha cleavage observed in cells. Furthermore, it is difficult to explain why Drosha cleavage fidelity on a given pri-miRNA can differ in different cells (Bofill-De Ros et al., 2019a; McCall et al., 2017) if it is determined only by these invariable features. Therefore, additional RNA elements that are amenable to regulation might contribute to Drosha cleavage fidelity.

By studying Drosha processing of the human pri-miR-9 family, we showed previously that the distorted and flexible structure of the pri-miR-9-1 lower stem promotes Drosha cleavage at an alternative site (Bofill-De Ros et al., 2019a). Pri-miR-9-2 and pri-miR-9-3, despite encoding the same mature miRNA, are cleaved by Drosha at a single site. As a result, a 5’ isomiR (miR-9-5p-alt) is exclusively generated from pri-miR-9-1 and regulates a distinctive set of target genes in low grade glioma. The case study of pri-miR-9 provided the first evidence linking pri-miRNA tertiary structure with Drosha cleavage fidelity. However, it remains unclear to what extent this applies to pri-miRNA processing in general.

Here, by analyzing *in vitro* Drosha cleavages of over 240,000 variants of pri-miR-16-1, pri-miR-30a and pri-miR-125a, we systematically investigated the relationship between pri-miRNA structure and Drosha cleavage fidelity. We report that structural flexibility introduced by unpaired regions and nearby G:U wobble pairs along the pri-miRNA stem leads to increased alternative cleavage of Drosha, which in turn impacts Dicer processing and hence contributes to 5’ isomiR production from both 5’ and 3’ arms of a pre-miRNA. Furthermore, we performed hypothesis-driven mutagenesis on pri-miR-9 and validated these conclusions on Drosha processing in cells. By analyzing data deposited in The Cancer Genome Atlas (TCGA), we provide evidence that alternative cleavage of pri-miRNAs is a tunable process that responds to the levels of pri-miRNA-associated RBPs. Together, our findings reveal that Drosha cleavage fidelity can be modulated by altering pri-miRNA structure, a mechanism by which cells might regulate 5’ isomiR biogenesis in tumors.

## RESULTS

### More flexible pri-miRNA structure correlates with alternative Drosha cleavages

To study how pri-miRNA structure impacts Drosha cleavage fidelity, we took advantage of a published dataset (Fang and Bartel, 2015) originally used to identify motifs required for efficient Drosha processing. Pri-miR-16-1, pri-miR-30a, pri-miR-125a and >240,000 variants with various sequences mutated along the pri-miRNA stem were cleaved by Drosha *in vitro*. By analyzing the sequences of the cleavage products, specifically the 5’ miRNA-offset RNA (5’ moR) (Berezikov et al., 2011), we observed on average ~150 cleavage events per variant and determined the corresponding Drosha cleavage site (Figure 1A). Consistent with previous reports (Berezikov et al., 2011; Bofill-De Ros et al., 2019a; Kwon et al., 2019; Ma et al., 2013; Shi et al., 2009), a portion of Drosha cleavages occurred at positions other than the canonical site for all three pri-miRNAs (Figure S1A), indicating that alternative Drosha processing is an intrinsic phenomenon.

We predicted the minimum free energy (MFE) of each pri-miR-16-1 variant and used it as an indication of the overall structural flexibility. Variants less structured (more flexible) than the wild-type pri-miR-16-1 (−37 kcal/mol) were processed by Drosha with an alternative cleavage frequency 2 to 6 times higher (Figure 1B), indicating that structural flexibility of pri-miRNA is inversely correlated with Drosha cleavage fidelity. Supporting this idea, pri-miR-16-1 variants with the highest alternative cleavage rates (top 1%) had the lowest average MFE value, while those with the lowest alternative cleavage rates (bottom 1%) had the highest average MFE value (Figure 1B). The same analysis with pri-miR-30a, pri-miR-125a and their variants generated similar results (Figure S1B–S1E), demonstrating that more flexible pri-miRNA structures were processed by Drosha with a lower cleavage fidelity.

Next, we grouped pri-miR-16-1 sequence variants (~80,000) based on their predicted secondary structures (N=1308 structural variants) with each group sharing the same structure. The average alternative Drosha cleavage ratio of each group was calculated and then compared to that of the group resembling wild-type pri-miR-16-1 structure (Figure 1C). Several groups showed increased alternative cleavage with extremely low p-values (<10^−150^), indicating that variants in these groups had a rather homogeneous change of alternative cleavage regardless of their sequence variations. It suggested that these particular structures, but not their sequences per se, underlie the increased alternative cleavage. We selected the top eight structural variants for further characterization (Figure 1D). Three of them (structures 1, 4 and 8) presented a disruption of the basal junction while variant 6 contained an unpaired region at the position of the mGHG motif, confirming the critical role of both basal junction (Ma et al., 2013) and mGHG motif (Fang and Bartel, 2015; Kwon et al., 2019) in determining Drosha cleavage site. We could not identify any consistent RNA secondary folding features among the other structural variants (2, 3, 5 and 7). Instead, after measuring their tertiary structure differences by the root-mean-square deviation (RMSD), we found that they all had relatively high variation compared to pri-miR-16-1 (WT) (Figure 1E). The same analysis on pri-miR-30a and pri-miR-125a datasets generated similar conclusions (Figure S1F and S1G). Consistent with the insights gained from the case study of pri-miR-9 (Bofill-De Ros et al., 2019a), these results indicate that overall distortion and flexibility of a pri-miRNA stem, a tertiary structure feature, determines to a large extent alternative Drosha cleavage.

### Unpaired internal loops lead to alternative Drosha cleavages

The four pri-miR-16-1 structural variants identified with a high ratio of alternative cleavage all contained a relatively large internal bulge along the stem (Figure 1D). To test whether the structural flexibility resulting from the internal loop leads to alternative Drosha cleavage, we further classified sequence variants within each structure according to the number of potential base-pairs formed in the internal loop (Figure 2A). For all structural variants tested, we observed reduced alternative Drosha cleavage when the internal loop could be partially paired (Figure 2B). The same results were obtained with a similar analysis of pri-miR-30a and pri-miR-125a (Figure S2A), suggesting that pri-miRNA tertiary structural flexibility associated with unpaired regions causes alternative Drosha cleavage *in vitro*.

**Figure 2.**
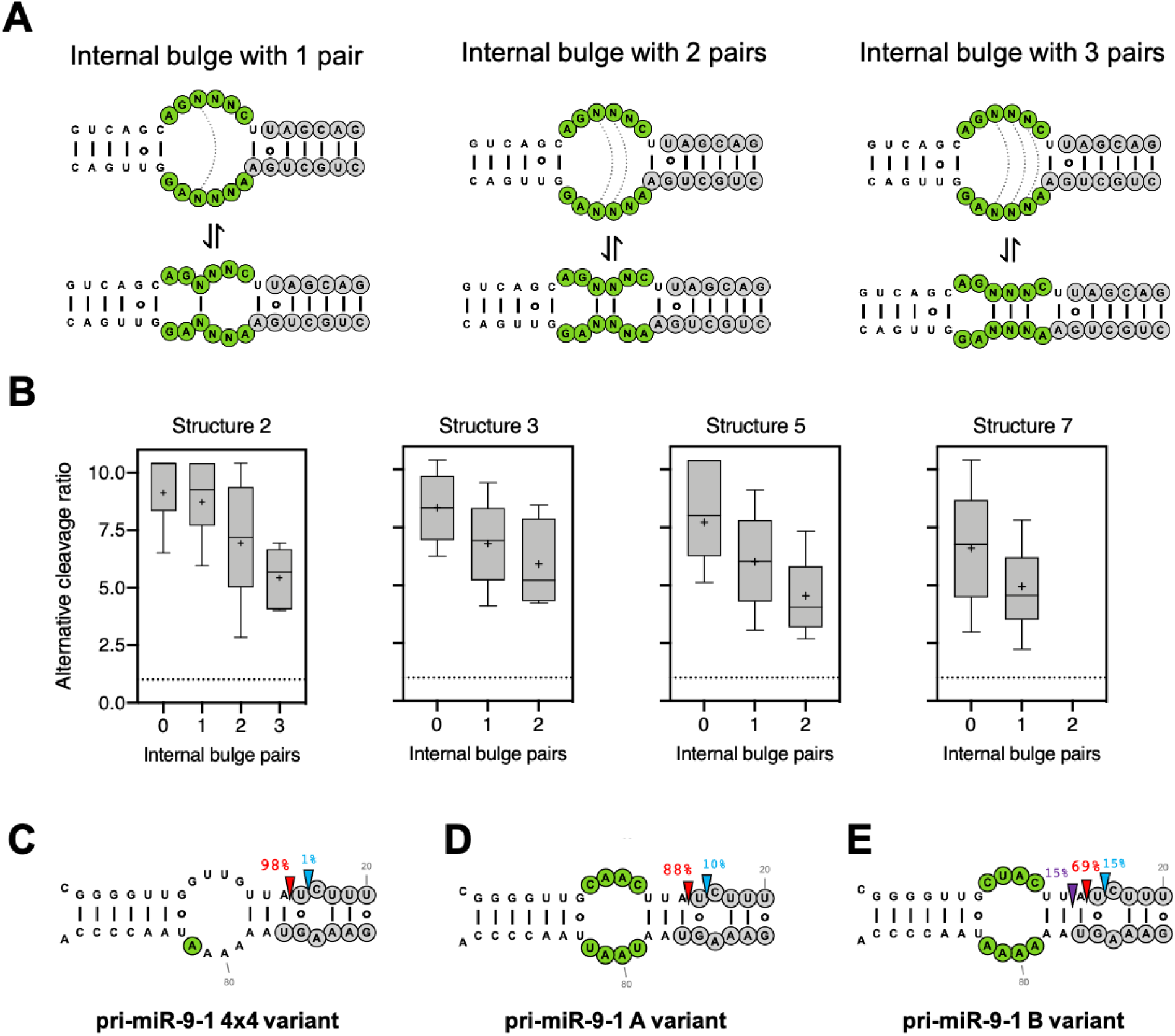
Unpaired internal loops lead to alternative Drosha cleavages. (A) Scheme of the internal bulge of pri-miR-16-1 structural variant 2 in equilibrium with structures that can close the bulge with one, two or three potential base-pairs. (B) Box plots with the alternative cleavage ratio of sequence variants that can form different numbers of base-pairs in the structural variant groups 2, 3, 5, and 7, previously described in Figure 1D. (C, D, E) HEK293T cells transfected with plasmids expressing pri-miR-9-1 lower stem variants. Nucleotides changed from the wild-type sequence are indicated in green. Small RNAs were subjected to deep sequencing. After being mapped to the corresponding pri-miR-9 structure, the percentage of sequences starting at a position relative to the total number of miR-9 reads was used to infer the Drosha site cleavage percentage. Canonical and alternative cleavage sites are indicated with a red, blue and purple arrow respectively.

To test this in living cells, we took advantage of pri-miR-9-1, the Drosha processing of which has been well-characterized: an asymmetrical internal bulge (4×3) at the lower stem of pri-miR-9-1 is responsible for its alternative Drosha cleavage (~15%) which can be reduced to <1% by correcting the asymmetry (Bofill-De Ros et al., 2019a). The resulting symmetrical bulge (4×4) can form two internal base-pairs (Figure S2B), suggesting this partial pair as the source of reduced alternative Drosha cleavage (Figure 2C). To test this, we disrupted the internal pairs by mutagenesis and measured their Drosha cleavage in HEK293T cells by deep sequencing. The resulting pri-miR-9-1 mutants, while maintaining the 4×4 symmetrical bulge, had greater flexibility (Figure S2C). As expected, we observed substantial amounts of alternative Drosha cleavage (Figure 2D and 2E). Together, these results demonstrate that unpaired bulges contribute to pri-miRNA structural flexibility, which in turn leads to alternative Drosha cleavage both *in vitro* and *in vivo*.

### G·U wobble pairs contribute to alternative Drosha cleavage by enhancing pri-miRNA structural flexibility

G·U pairs are known to disrupt dsRNA A-form helix structure (Varani and McClain, 2000). Therefore, we sought to investigate how they contribute to structure-flexibility-mediated alternative Drosha cleavage. To this end, we analyzed all pri-miR-16-1 sequence variants that are predicted to have the same secondary structure but different numbers of G·U pairs compared to the wild-type pri-miR-16-1. Compared to variants having the same number of G·U pairs as the wild-type (N=1,948 sequence variants), those with additional G·U pairs (N=2,436 sequence variants) were processed by Drosha with a higher average alternative cleavage ratio (Figure 3A). The same analysis for pri-miR-30a and pri-miR-125a variants showed a similar result (Figure S3), indicating that G·U pairs promote alternative Drosha cleavage. However, the average effect was subtle with a large variation, suggesting that not all G·U pairs contribute equally. We analyzed the impact of the G·U position on alternative cleavage and found that changes at certain positions had a much larger effect (Figure 3B). The positions of these hotspots were not consistent among pri-miR-16-1, pri-miR-30a and pri-miR-125a, suggesting that the effect is unlikely caused by specific interactions between Drosha and pri-miRNA substrate. Instead, incorporating G·U pairs at positions near existing bulges increased the alternative cleavage rate (Figure 3B–D), suggesting that G:U pairs promote alternative Drosha cleavage by enhancing existing structural flexibility.

**Figure 3.**
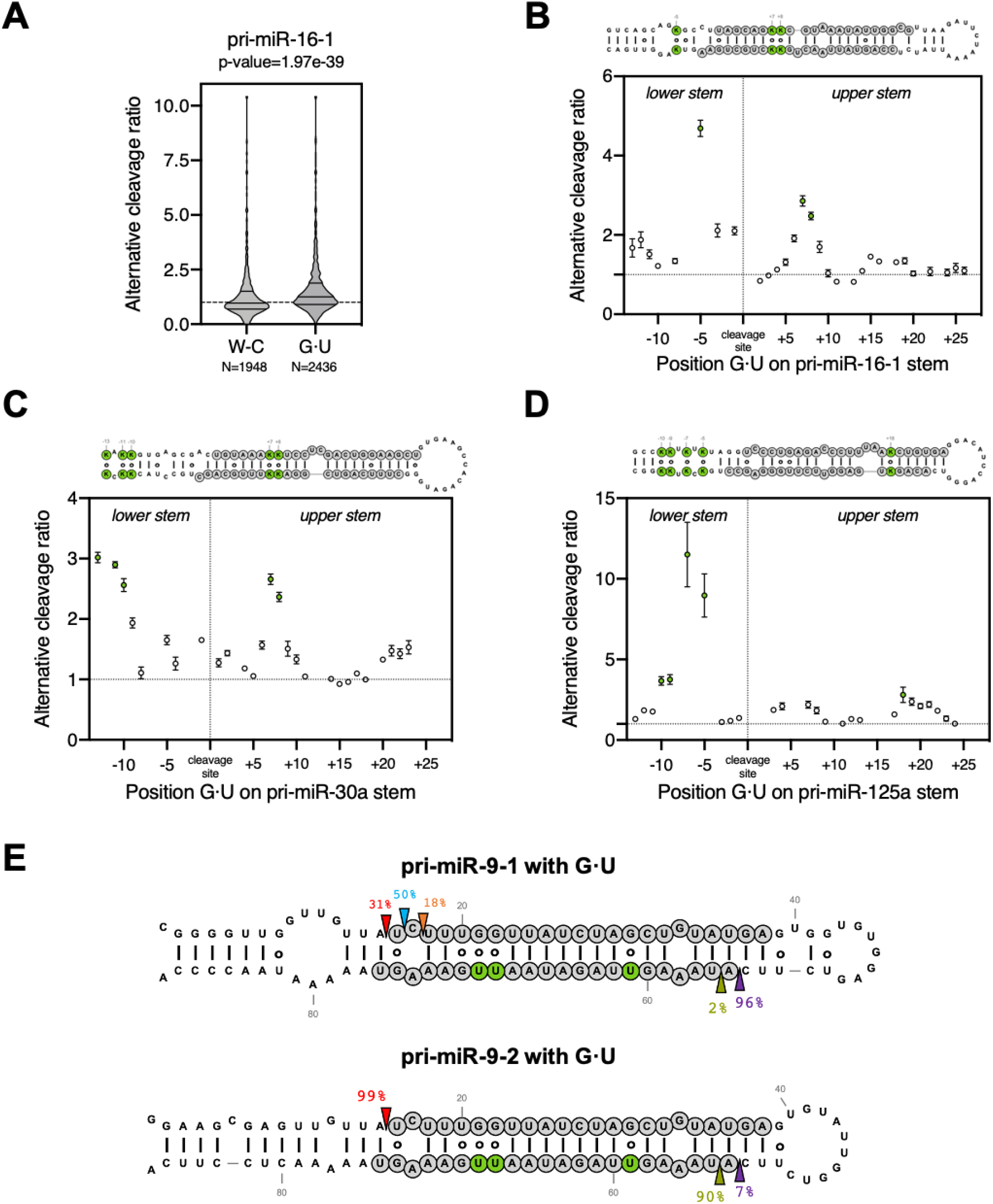
G·U wobble pairs contribute to alternative Drosha cleavage by enhancing pri-miRNA structural flexibility. (A) Violin plot of the alternative cleavage ratio of pri-miR-16-1 variants with wild-type native structure. W-C indicate variants with the same number of G·U pairs as the wild-type, and G·U indicates variants with additional G·U pairs. (B, C, D) Analysis of the effect of the addition of single G·U wobble pairs on alternative cleavage based on their position in the stem of pri-miR-16-1, pri-miR-30a and pri-miR-125a (dots indicate mean and standard error). Secondary structures indicate in green the location of G·U wobble pairs with the largest impact on alternative Drosha cleavage. (E) HEK293T cells transfected with plasmids expressing either pri-miR-9-1 or pri-miR-9-2 variants with additional G·U wobble pairs (indicated in green). Small RNAs mapping to each pri-miRNA were used to infer the percentage of Drosha and Dicer cleavage at each position. Canonical cleavage sites of Drosha and Dicer are indicated with red and purple arrows, while blue, orange and green indicated alternative cleavage sites.

To test this idea, we took advantage of pri-miR-9-1 and pri-miR-9-2, with the former having a distorted tertiary structure and the latter being structurally rigid (Bofill-De Ros et al., 2019a). Replacing multiple Watson-Crick G-C pairs by G·U pairs along the stem significantly increased the alternative cleavages of pri-miR-9-1 but had no effect on pri-miR-9-2 when expressed in HEK293T cells (Figure 3E). Together, these results demonstrate that a G:U pair per se has minimal impact on Drosha cleavage. However, G:U wobble pairs contribute to alternative Drosha cleavage by enhancing structural flexibility of pri-miRNAs.

### Alternative Drosha cleavage results in 5’ isomiRs from both strands of pre-miRNAs

Drosha and Dicer alternative cleavages generate 5’ isomiRs from the 5p arm and 3p arm of pre-miRNAs respectively (Figure S4A). Given that Drosha processing determines to a large extent where Dicer cuts (Macrae et al., 2006; Zhang et al., 2004), alternative Drosha cleavage may also contribute to the production of 3p isomiRs. To test this, we expressed pri-miR-9-1 and pri-miR-9-2 separately in HEK293T cells and examined the biogenesis of their 3p isomiRs. Northern blot analysis revealed that both pri-miR-9 transcripts produced 3p isomiRs (miR-9-3p) (Figure 4A). By deep sequencing, we found that these miR-9-3p reads were composed of two populations: a canonical miR-9-3p (miR-9-3p-can) as annotated in the miRBase and a 5’ isomiR (miR-9-3p-alt) that begins one nucleotide downstream (Figure 4B). Interestingly, their relative abundances differed between pri-miR-9-1 and pri-miR-9-2. While miR-9-3p-can was the dominant isomiR processed from pri-miR-9-1 (~70%), it only accounted for ~30% of reads generated from the pri-miR-9-2 (Figure 4B). These results indicate that the dominant Dicer cleavage site on pri-miR-9-1 is one nucleotide upstream of the Dicer cleavage site on pri-miR-9-2. Because the current model indicates that Dicer cuts at a fixed distance from where Drosha cuts (Macrae et al., 2006; Park et al., 2011; Zhu et al., 2018), this shift of Dicer cleavage position aligns well with the alteration of Drosha cleavage sites (Figure 4B). This suggests that the decreased amount of miR-9-3p-alt generated from pri-miR-9-1 is likely a result of its alternative Drosha cleavage. To test this, we examined the biogenesis of 3p isomiRs in multiple pri-miR-9-1 mutants, all of which have the same upper stem and loop sequence but different Drosha cleavage patterns due to variation at the lower stem sequence (Figure 4C and S4B). As expected, reduced alternative Drosha cleavage led to higher levels of miR-9-3p-alt whereas increased alternative Drosha cleavage resulted in lower levels of miR-9-3p-alt (Figure 4D). Together, these results demonstrate that Drosha processing contributes to the production of 3p isomiRs by impacting Dicer cleavage.

**Figure 4.**
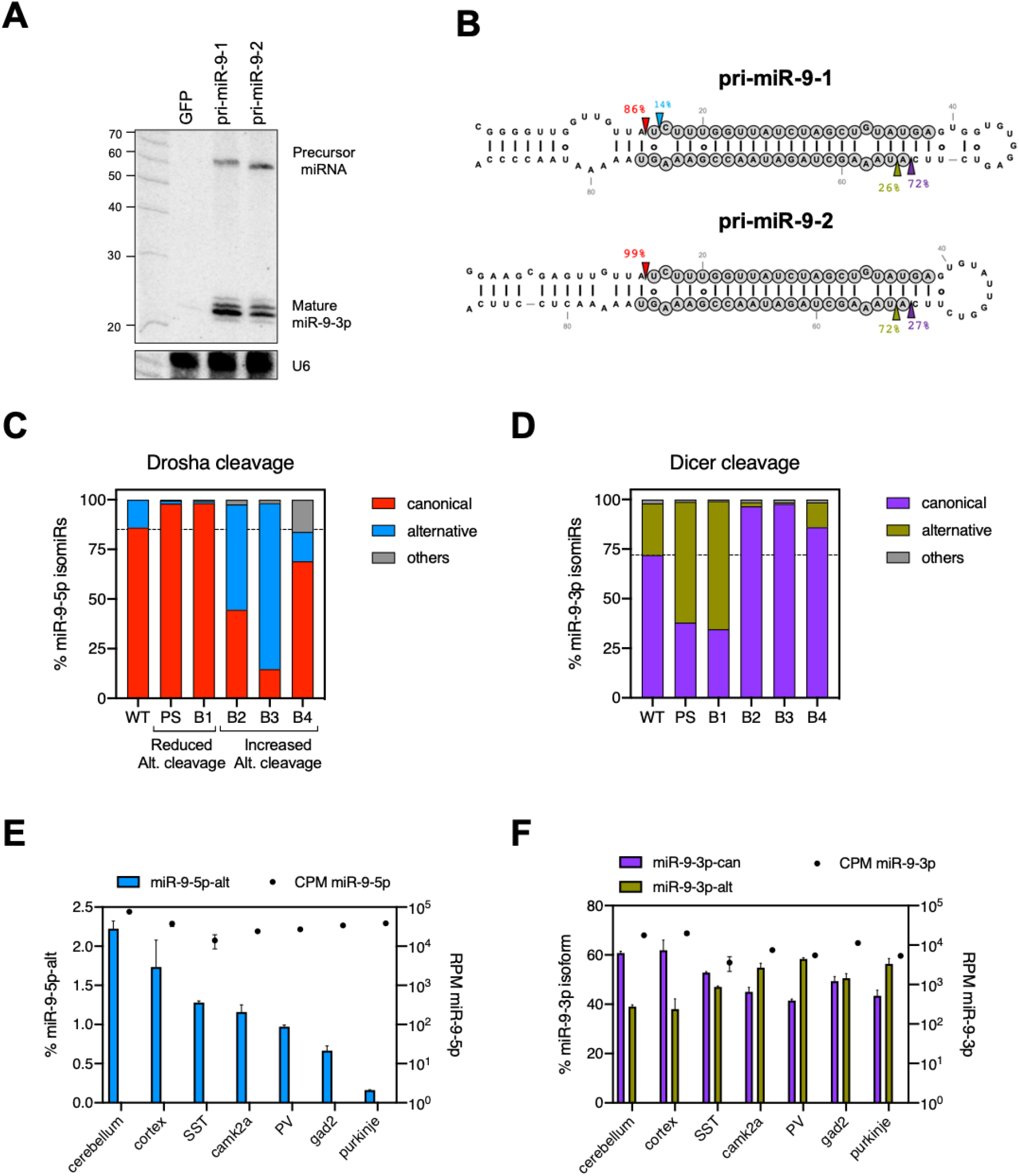
Alternative Drosha cleavage results in 5’ isomiRs from both strands of pre-miRNAs. (A) HEK293T cells transfected with plasmids expressing either pri-miR-9-1 or pri-miR-9-2 were subjected to northern blotting to detect miR-9-3p expression. (B) Small RNAs mapping to each pri-miRNA paralog were used to infer the percentage of Drosha and Dicer cleavage at each site. (C, D) Frequency of reads with the canonical or alternative 5’ end for miR-9-5p (Drosha cleavage) and miR-9-3p (Dicer cleavage) for different pri-miR-9-1 variants on the lower stem. Canonical and alternative 5′ends were defined based on miRBase annotation for the mature miRNA (Kozomara et al., 2019). (E, F) Small RNA deep sequencing data from different mouse brain tissues (cerebellum, cortex) and cell-types (Purkinje cells, Camk2α cells, parvalbumin (PV) neurons and neuropeptide somatostatin (SST) neurons) were re-analyzed to assess miR-9-5p and miR-9-3p isoforms (left axis) and overall expression (RPM, counts per million, right axis).

To extend our conclusion beyond cultured cells, we examined miR-9 biogenesis in mouse tissues. Consistent with previous studies, miR-9 expression was highly specific to the brain (Figure S4C). Analyses of a previously published sRNA-seq dataset (He et al., 2012) revealed that miR-9 is expressed in multiple brain tissues and various neuronal cells. Despite the rather homogenous expression level (~40,000 CPM), the percentage of the 5p isomiR (miR-9-5p-alt) ranged from 2.2% in cerebellum to less than 0.2% in Purkinje cells (Figure 4E), suggesting variations in alternative Drosha cleavages. The distribution of 3p isomiRs was changed accordingly: a higher percentages of miR-9-3p-can was observed in tissues/cells with a higher percentage of miR-9-5p-alt; while more miR-9-3p-alt was observed in those with a lower percentage of miR-9-5p-alt (Figure 4F), which is consistent with the pattern observed in HEK293T cells. These results suggest that alternative Drosha cleavage could play a biological role in regulating 5’ isomiR biogenesis.

### Alternative Drosha cleavage is subjected to cellular regulation

Drosha cleavage fidelity on a given pri-miRNA varies among cell types (Bofill-De Ros et al., 2019a; McCall et al., 2017), suggesting cellular regulation. To further test this idea, we took advantage of The Cancer Genome Atlas (TCGA), where a large amount of miRNA-seq and corresponding RNA-seq data are available. We measured the relative levels of 5’ isomiRs for the top 200 abundant miRNAs in 1015 Breast Invasive Carcinoma (BRCA) samples. Comparing these to normal tissue controls (N=104), we observed a subtle but significant increase in 5’ isomiRs. We made a similar observation was made with samples obtained from Kidney Renal Clear Cell Carcinoma (KIRC) and Uterine Corpus Endometrial Carcinoma (UCEC) patients, indicating that the upregulation of 5’ isomiRs is not limited to one type of cancer (Figure 5A). These results suggest that Drosha processing is altered during tumorigenesis, resulting in 5’ isomiRs that could potentially impact tumor progression. Interestingly, pri-miRNAs cleaved by Drosha with high fidelity in normal cells were more resistant to such an alteration whereas pri-miRNAs with low Drosha cleavage fidelity were more prone to the increase in alternative Drosha cleavage (Figure 5B).

**Figure 5.**
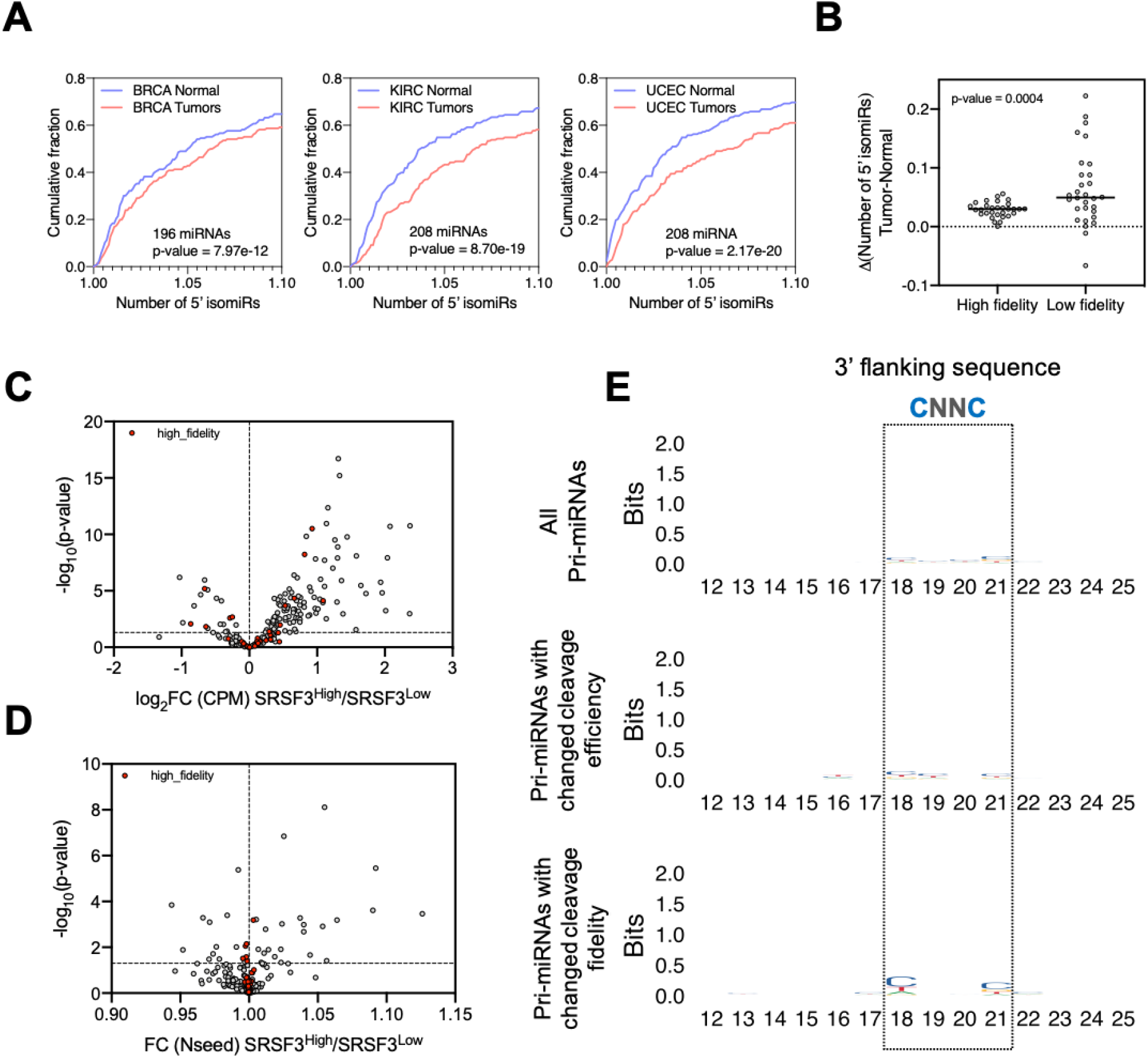
Alternative Drosha cleavage is subjected to cellular regulation. (A) The relative number of 5’ isomiRs was measured using an inverted Simpson diversity index. Using this measurement, we analyzed all highly expressed miRNAs comparing normal and primary tumor samples from BRCA, KIRC and UCEC. Results are plotted as a cumulative distribution for all miRNAs between normal and tumoral samples. (B) Delta of the number of 5’ isomiRs between BRCA primary tumors and normal samples. High fidelity and low fidelity groups were defined as the top and bottom 30 miRNAs, out of the top 100 most expressed, according to their levels of 5’ isomiRs in normal samples. (C) Volcano plot of the changes in mature miRNA expression between the two sets of BRCA primary tumor samples with high and low levels of SRSF3 expression (N=100 samples/group). The group of miRNAs processed with high fidelity (defined in Figure 5B) is shown in red. (D) Similarly, volcano plot of the changes in mature miRNA Number of 5’ isomiRs (Nseed) between BRCA primary tumors with high/low levels of SRSF3. (E) Motif logo of the 3’ flanking sequence of the following groups: all pri-miRNAs included in the analysis (upper panel), pri-miRNAs with differential cleavage efficiency (miRNA expression) (FC>1 and p<0.05) (middle panel), pri-miRNAs with differential cleavage fidelity (p<0.05) (lower panel). Box indicates the position for binding of SRSF3 previously described.

Given the role that structure plays in defining Drosha cleavage sites, the regulation of 5’ isomiR production could be achieved, at least in part, via modulating pri-miRNA structures by association of RBPs. To test this, we sought to investigate whether SRSF3, an RBP known to bind to pri-miRNAs at a “CNNC” motif downstream of the Drosha cleavage site, could impact alternative Drosha cleavages. To this end, we compared the miRNA profiles between BRCA patients with high levels of SRSF3 (top 10%, 100 samples) and relatively low levels of SRSF3 (bottom 10%, 100 samples). Consistent with the known role of SRSF3 in promoting miRNA biogenesis (Auyeung et al., 2013; Kim et al., 2018), expression levels of most miRNAs were higher in samples with a higher level of SRSF3, confirming our approach (Figure 5C). The fidelity of Drosha cleavage on a subset of pri-miRNAs also varied between the two groups (Figure 5D). A “CNNC” motif was enriched in this subset of pri-miRNAs, indicating that the observed changes in Drosha fidelity were likely a direct effect of SRSF3 association (Figure 5E).

Similar analyses of a set of RBPs known to associate with pri-miRNAs (Nussbacher and Yeo, 2018; Treiber et al., 2017) generated similar results, whereas the level of randomly selected RBP RBM22 as well as a non-RBP Tubulin (TUBA1A), had marginal, if any, effects on 5’ isomiR profiles (Figure S5A). Different RBPs had different impacts: while a higher level of DDX3X, DDX21, hnRNPA1 or hnRNPH2 generally promoted alternative Drosha cleavage (Figure S5B), FUS and hnRNPH1 apparently prevented Drosha alternative processing (Figure S5C). In all cases, pri-miRNAs with no change in Drosha cleavage fidelity during tumorigenesis also showed no impact from these RBPs, indicating that RBP-mediated regulation of alternative Drosha cleavage is limited to those pri-miRNAs without a well-defined Drosha cleavage site.

## DISCUSSION

As the initial step licensing miRNA production, Drosha cleavage of pri-miRNAs has been extensively studied. A comprehensive set of structural and sequence features of pri-miRNAs have been identified to play an important role in determining Drosha cleavage efficiency and fidelity. However, most endogenous pri-miRNAs have only a subset of these features, suggesting certain evolutionary advantages to having non-optimal processing. Here, by analyzing tens of thousands of *in vitro* Drosha cleavage events, we provide robust statistical evidence indicating that pri-miRNA structural flexibility introduced by internal loops and G:U wobbles is positively correlated with alternative Drosha cleavage. Using mutagenesis study of pri-miR-9, we established causality and validated these conclusions in living cells. Pri-miRNA structural flexibility, different from previously identified features, is amenable to cellular regulation. Indeed, we provided evidence that the level of a set of pri-miRNA binding proteins, including SRSF3, correlates with the use of alternative Drosha cleavage sites in tumors. Given the prevalence of internal bulges and G:U wobbles along the pri-miRNA stem, our findings support a model in which Drosha cleavage fidelity is regulated by modulating pri-miRNA structure via association of RBPs.

It is intriguing to speculate why pri-miRNAs with distorted stems are processed by Drosha with a lower cleavage fidelity. It is possible that the higher flexibility of pri-miRNAs enables them to fold into several distinct suboptimal structures when complexing with Drosha. In this case, the various cleavage sites may be a result of different configurations of the catalytic center and substrate. Indeed, the flexible lower stem of pri-miR-9-1, which has a ~15% chance to be cut by Drosha at an alternative site, can fold into two suboptimal structures (Figure S6A). Two constructs (SUB1 and SUB2) designed to mimic these suboptimal conformations were processed by Drosha differently: Drosha cut SUB1 primarily at the canonical site (Figure S6B) whereas SUB2 was processed with a higher level of miscleavage than the wild-type structure (Figure S6C). This suggests that the overall Drosha cleavage profile of pri-miR-9-1 may be an ensemble of different configurations between Drosha and two suboptimal folds of pri-miR-9-1. Future high-resolution structures of the ternary complex formed by Drosha, DGCR8 and pri-miRNA should give additional insights (Jin et al., 2020; Partin et al., 2020).

By chemical probing approaches, a recent report measured the endogenous RNA structures on a genome-wide scale (Sun et al., 2019). When comparing RNAs extracted from different cellular compartments, the authors find that RNA structures vary less *in vitro* than *in vivo*. In particular, conserved pri-miRNAs form relatively stable structures *in vitro* yet are highly influenced by cellular factors, resulting in different RNA-folds *in vivo*. A follow-up study demonstrated that these distinct structures of pre-miRNAs correlate with Dicer cleavage efficiency and fidelity (Luo et al., 2021). Furthermore, another study compared pri-miRNA processing *in vitro* and *in vivo* and found that cleavage fidelity is regulated *in vivo* by RBPs such as SRSF3 (Kim et al., 2021). These results further support our model that pri-miRNA structure is modulated to fine-tune 5’ isomiR production. We have previously shown that a Drosha isoform lacking the nuclear localization signal due to alternative splicing can process a subset of pri-miRNAs in the cytoplasm (Dai et al., 2016). It is possible that the cytoplasmic pri-miRNAs are processed with different Drosha cleavage fidelity due to changes in their structures. This might help to explain why up-regulation of cytoplasmic Drosha coincides with misregulation of miRNAs in tumors.

5’ isomiRs resulting from alternative Drosha cleavages play diverse biological roles. In particular, we have shown that 5’ isomiRs processed from pri-miR-9-1 regulates a distinct set of target mRNAs in low grade gliomas (Bofill-De Ros et al., 2019a). Here, we found that Drosha cleavage fidelity decreases in multiple cancers, resulting in numerous miRNAs with altered 5’ ends and seed sequences. This led us to hypothesize that Drosha processing is altered during tumorigenesis to produce 5’ isomiRs that impact tumor progression. Our finding that pri-miRNA-associated RBPs promote or inhibit 5’ isomiR production by modulating pri-miRNA structure provides one possible underlying mechanism. It is possible that only a subset of the aberrant 5’ isomiRs found in tumors have a functional role. Future studies identifying oncogenic or tumor suppressive 5’ isomiRs will provide additional insights.

## STAR⋆METHODS

### KEY RESOURCES TABLE

| REAGENT or RESOURCE | SOURCE | IDENTIFIER |
| --- | --- | --- |
| **Experimental Models: Cell Lines** |  |  |
| Human: 293T cells | ATCC | Cat# CRL-3216 |
| **Chemicals, Peptides, and Recombinant Proteins** |  |  |
| DMEM High glucose | GIBCO | Cat# 11965092 |
| Penicillin-Streptomycin (10,000 U/mL) | GIBCO | Cat# 15140163 |
| HyClone™ Fetal Bovine Serum | GE Healthcare | Cat# SH30070 |
| PolyJet™ DNA Transfection Reagent | SignaGen | Cat# SL100688 |
| PerfectHyb™ Hybridization Buffer | Sigma | Cat# H7033-1L |
| NEBNext Small RNA Library Prep | NEB | Cat# E7330L |
| Qubit™ dsDNA HS Assay Kit | ThermoFisher | Cat# Q32854 |
| Agilent High Sensitivity DNA kit | Agilent | Cat# 5067-4626 |
| MiSeq Reagent Kit v3 (150-cycle) | Illumina | Cat# MS1023001 |
| **Deposited Data** |  |  |
| FASTQ files | This paper | GEO: GSE172446 |
| **Other** |  |  |
| Previous miR-9 processing | (Bofill-De Ros et al., 2019a) | GEO: GSE108893 |
| Synthetic pri-miRNA libraries | (Fang and Bartel, 2015) | GEO: GSE67937 |
| Ago2-IP in mouse brain | (He et al., 2012) | GEO: GSE30286 |

### CONTACT FOR REAGENT AND RESOURCE SHARING

Further information and requests for resources should be directed to Shuo Gu.

## METHOD DETAILS

### Cell lines

HEK293T cells were maintained in DMEM high glucose (Gibco) supplemented with 10% heat-inactivated fetal bovine serum (Hyclone), 100 U/ml penicillin-streptomycin (Gibco) at 37°C. Cells were tested to be free of mycoplasma contamination. Transfections were performed using PolyJet™ DNA Transfection Reagent (SignaGen) according to the manufacturer’s instructions.

### Northern blot

Total RNA was isolated using Trizol (Life Technologies) and separated in denaturing gels, 20% (w/v) acrylamide with 8 M urea. RNA was transferred to Hybond-N1 membranes (Amersham Pharmacia Biotech), crosslinked and blocked (PerfectHyb™ Plus Hybridization Buffer - Sigma). MicroRNAs were detected using ^32^P-labeled oligonucleotide probes (miR-9-5p: GACTCATACAGCTAGATAACCAAAG, miR-9-3p: GACTTTCGGTTATCTAGCTTTAT). Images were obtained and analyzed using Amersham Typhoon (GE Healthcare).

### Small RNA NGS library preparation

Total RNA (5 μg) was ligated to the RNA 3’ adaptor using T4 RNA Ligase 2 - truncated (NEB), in the presence of RNase Inhibitor (NEB). RNA 5’ adaptor was ligated using T4 RNA Ligase 1 - high concentration (NEB) and 10 mM ATP. Ligated small RNAs were reverse transcribed using SuperScript® IV Reverse Transcriptase (Thermo-Fisher). Small RNA library cDNA was amplified and indexed using Phusion® High-Fidelity DNA polymerase (NEB). Constructs were purified in a 6% (w/v) native acrylamide gel based on the expected product size and purified by ethanol precipitation. Library quality was assessed using Qubit dsDNA HS Assay Kit (ThermoFisher) and Agilent High Sensitivity DNA kit (Agilent). Libraries were mixed together and prepared at a final concentration of 12 pM and run on MiSeq Reagent Kit v3 (Illumina) according to the manufacturer’s specifications.

### MicroRNA expression and 5’ isomiR analysis

The primary analysis of miRNA expression and 5’ isomiR analysis were performed using QuagmiR on the NCI Cancer Genomics Cloud (Bofill-De Ros et al., 2019b). The number of 5’ isomiRs was calculated using an inverse Simpson diversity index (Bofill-De Ros et al., 2021). This number measures the evenness of the 5’ isomiRs generated from each individual matfure miRNA arm. Normal tissue and primary tumor samples from TCGA were used to calculate the number of 5’ isomiRs corresponding to each tumor type. Based on their average expression levels, we selected the most abundant miRNAs (with at least an average of 1 CPM). To analyze the impact of different RBPs on miRNA processing, we ranked the cohort of BRCA primary tumors according to their FPKM for the selected gene. Two groups of one hundred samples with the highest and the lowest expression levels were generated. More details of the R analysis are reported on GitHub (https://github.com/Gu-Lab-RBL-NCI/Drosha-alternative-cleavage).

### Analysis of Microprocessor cleavage fidelity in a library of variants

Data from synthetic libraries of different pri-miRNA (pri-miR-16-1, pri-miR-30a and pri-miR-125a) was downloaded from GEO accession GSE67937. First, barcoded cleaved fragments data (barcode_cleavage_fragments.txt) were used as 5’ miRNA-offset RNA (5’ moR) to establish the frequency of Drosha cleavage at the canonical and alternative sites. Second, for each barcoded variant in the library (dictionary.txt), the expected minimum free energy and secondary structure were calculated using the RNAfold from the Vienna package (Gruber et al., 2008). Subsequent analyses of secondary structures were performed by aggregation of variants of alternative cleavage based on their corresponding dot-bracket notation. More details of the R analysis are reported on GitHub (https://github.com/Gu-Lab-RBL-NCI/Drosha-alternative-cleavage).

### Data availability

Small RNA-seq data is deposited at GEO with the accession number GSE172446 [Reviewer access: ijobkaeshbmtfcd] and GSE108893. As well as, previously published Ago2-IP in mouse brain (GSE30286). The Cancer Genome Atlas (TCGA) can be accessed via dbGaP study accession: phs000178.v11.p8.

### Statistical analysis

p-values were calculated using the Wilcoxon test, as indicated. p-value < 0.05 was considered statistically significant. Statistical analysis was performed in RStudio and GraphPad Prism7 statistical software.

