## Supplemental information for "Flexible pri-miRNA structures enable tunable production of 5’ isomiRs"

[3]Lead contact

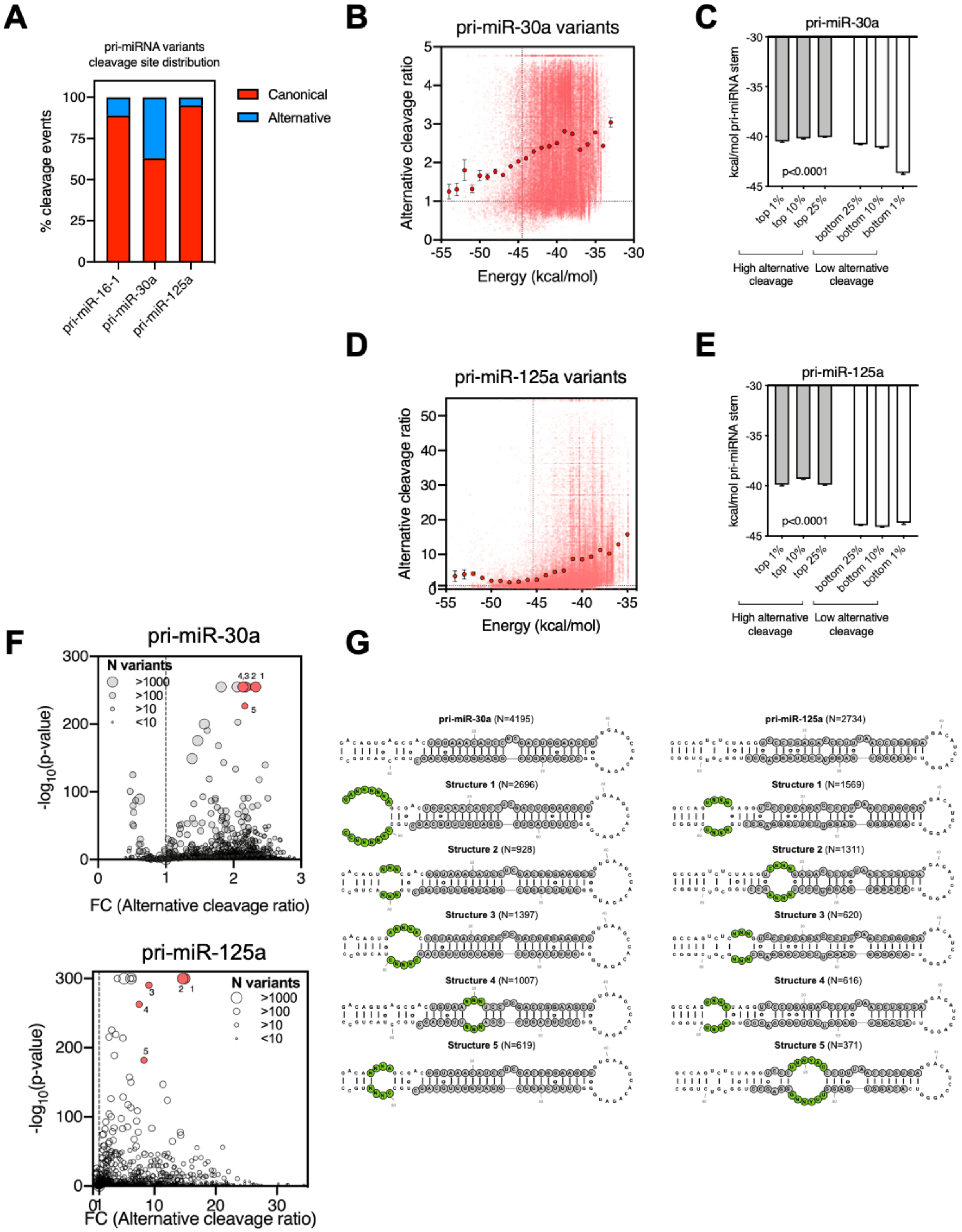
A
pri-miRNA variants cleavage site distribution
% cleavage events
Canonical
Alternative
pri-miR-16-1
pri-miR-30a
pri-miR-125a
B
pri-miR-30a variants
Alternative cleavage ratio
Energy (kcal/mol)
C
pri-miR-30a
kcal/mol pri-miRNA stem
p<0.0001
top 1%
top 10%
top 25%
bottom 25%
bottom 10%
bottom 1%
High alternative cleavage
Low alternative cleavage
D
pri-miR-125a variants
Alternative cleavage ratio
Energy (kcal/mol)
E
pri-miR-125a
kcal/mol pri-miRNA stem
p<0.0001
High alternative cleavage
Low alternative cleavage
F
pri-miR-30a
N variants
>1000
>100
>10
<10
-log10(p-value)
FC (Alternative cleavage ratio)
pri-miR-125a
N variants
-log10(p-value)
FC (Alternative cleavage ratio)
G
pri-miR-30a (N=4195)
pri-miR-125a (N=2734)
Structure 1 (N=2696)
Structure 1 (N=1569)
Structure 2 (N=928)
Structure 2 (N=1311)
Structure 3 (N=1397)
Structure 3 (N=620)
Structure 4 (N=1007)
Structure 4 (N=616)
Structure 5 (N=619)
Structure 5 (N=371)

**Figure S1. More flexible pri-miRNA structure correlates with alternative Drosha cleavages**

(A) Percentage of Canonical and Alternative Drosha cleavage events observed in the *in vitro* processing of pri-miR-16-1, pri-miR-30a and pri-miR-125a variant libraries. (B, D) Scatter plot of the alternative cleavage ratio (relative to pri-miR-30a and pri-miR-125a wild-type sequence, respectively) as a function of the minimum folding energy (kcal/mol) of each sequence variant. (C, E) Minimum folding energy (kcal/mol) of pri-miR-30a and pri-miR-125a sequence variants with high and low alternative cleavage levels. (F) Average alternative cleavage ratios of pri-miR-30a and pri-miR-125a sequence variants with the same predicted secondary structure were plotted against the p-values which were calculated by comparison to the variants with the same secondary structure as their native wild-type pri-miRNA structure (N=4,195 and 2,734, respectively) using Wilcoxon test. The size of the dots indicate the number of sequence variants supporting that structure group. (G) Structural variants with the most significant increases in alternative Drosha cleavage compared to variants with the native structure. N indicates the number of sequence variants supporting that structure group. Nucleotides involved in the structural feature are indicated in green and shown in the IUPAC nucleotide code.

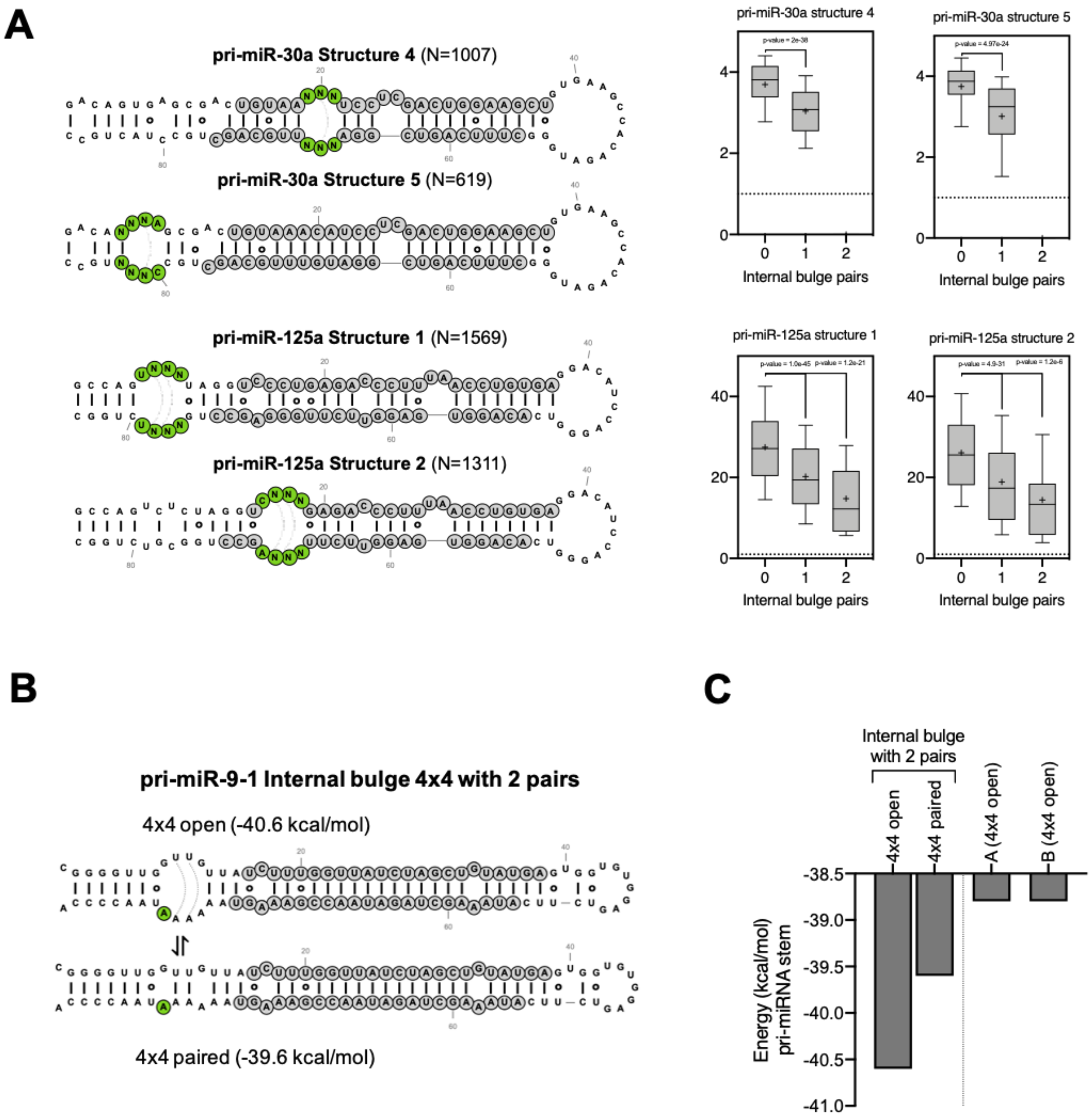


**Figure S2. Unpaired internal loops lead to alternative Drosha cleavages**

(A) Left, scheme of the internal bulges of pri-miR-30a and pri-miR-125a structural variant groups analyzed. Right, box plots with the alternative cleavage ratio of sequence variants that can form different numbers of base-pairs in the structural variant groups. (B) Scheme of the pri-miR-9-1 4x4 variant in the conformations of open and paired internal loop. (C) Predicted energy associated with the distinct potential pairings of pri-miR-9-1 variants 4x4, A and B, analyzed in Figures 2C–D and 2E.

A

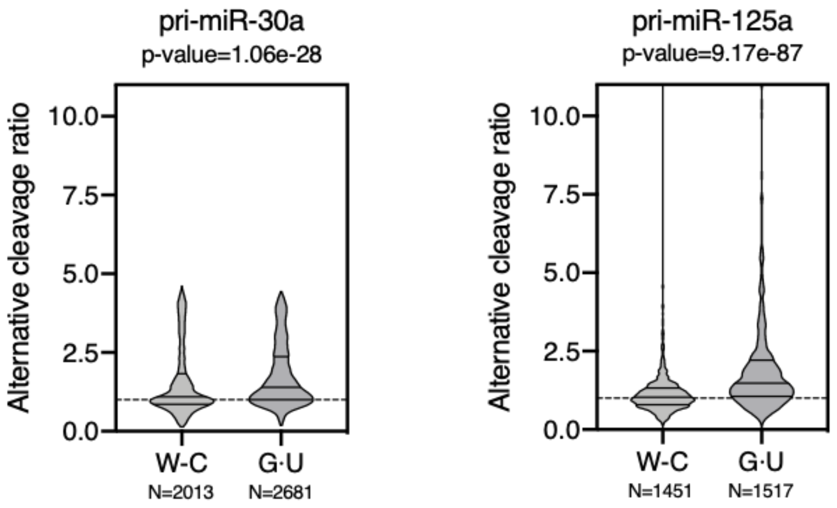


**Figure S3. G·U wobble pairs contribute to alternative Drosha cleavage by enhancing pri-miRNA structural flexibility**

(A) Violin plot of the alternative cleavage ratio of pri-miR-30a and pri-miR-125a variants with wild-type native structure. W-C indicate variants with the same number of G·U pairs as the wild-type, and G·U indicates variants with additional G·U pairs. N indicates the number of sequence variants supporting each classification.

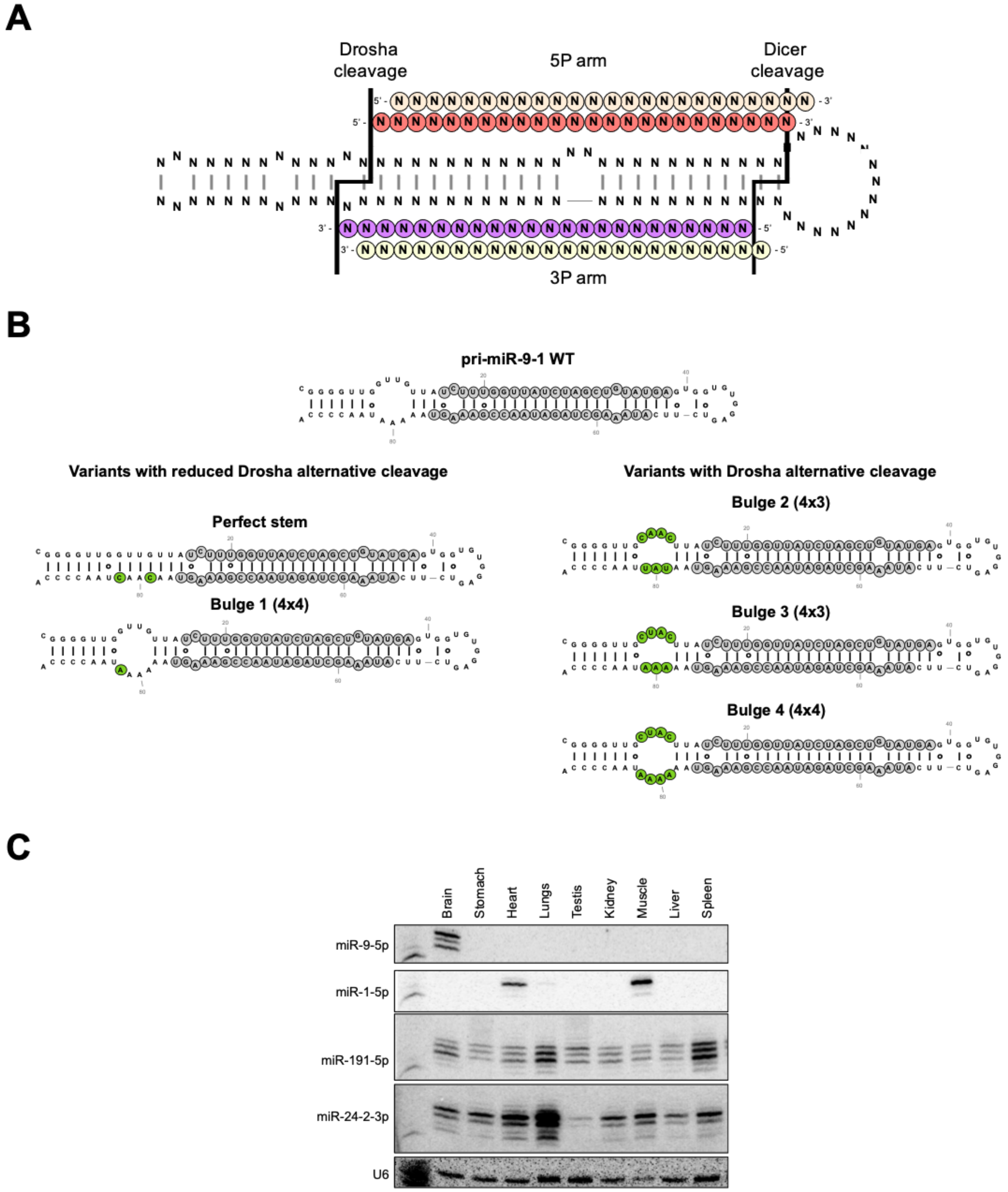
A
Drosha cleavage
5P arm
Dicer cleavage
3P arm
B
pri-miR-9-1 WT
Variants with reduced Drosha alternative cleavage
Perfect stem
Bulge 1 (4x4)
Variants with Drosha alternative cleavage
Bulge 2 (4x3)
Bulge 3 (4x3)
Bulge 4 (4x4)
C
Brain
Stomach
Heart
Lungs
Testis
Kidney
Muscle
Liver
Spleen
miR-9-5p
miR-1-5p
miR-191-5p
miR-24-2-3p
U6

**Figure S4. Alternative Drosha cleavage results in 5' isomiRs from both strands of pre-miRNAs**

(A) Scheme of pri-miRNA processing. Solid lines indicate the 5' ends generated by Drosha and Dicer cleavage. In light orange and yellow, illustrative examples of 5' isomiRs generated by the alternative cleavage of both enzymes. (B) Scheme of the pri-miR-9-1 wild-type and variants analyzed in Figures 4C–D. Nucleotides changed from the wild-type sequence are indicated in green. (C) RNA extracted from wild-type mice tissues were subject to northern blotting to detect expression of different miRNAs.

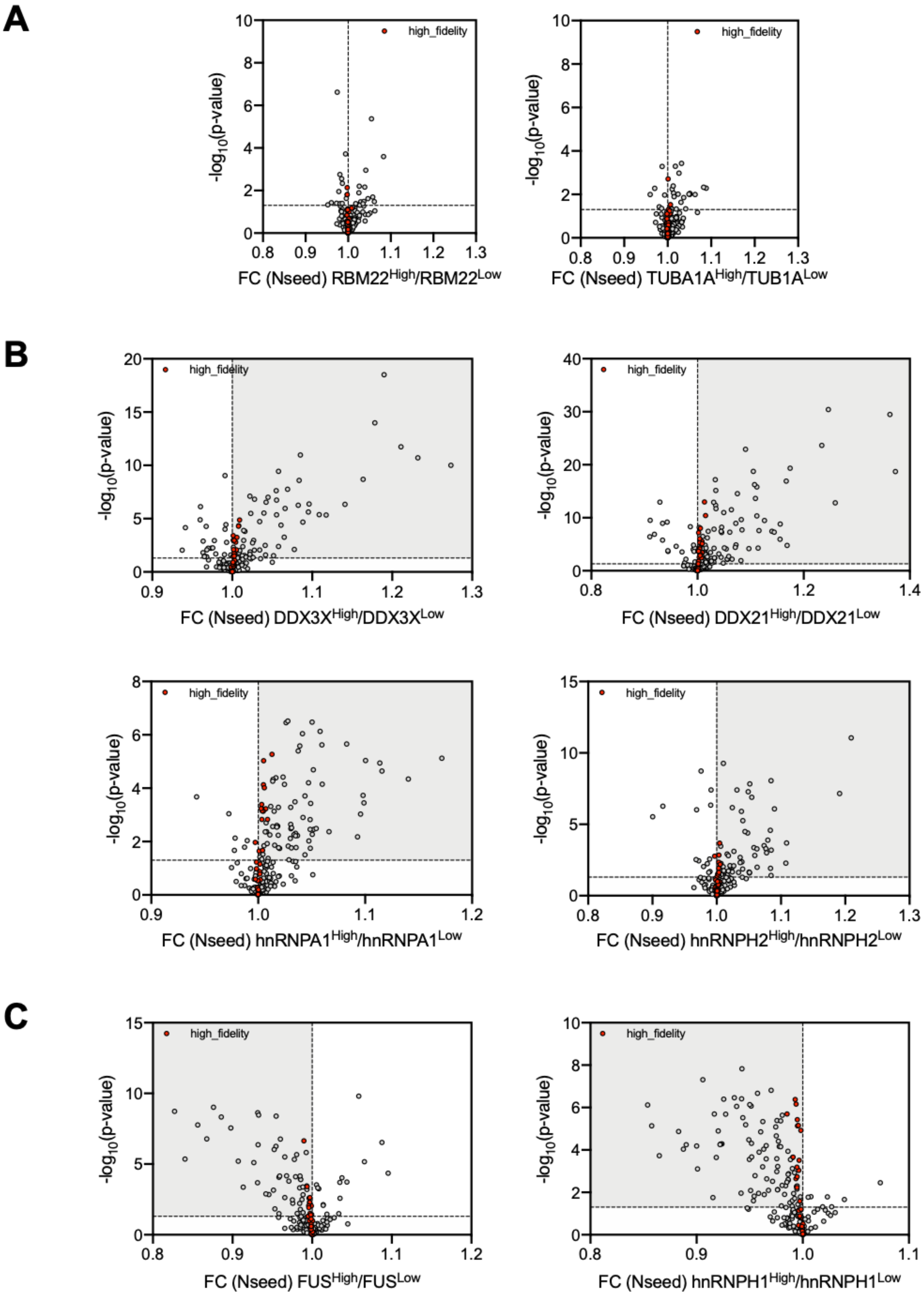
A
high_fidelity
-log10(p-value)
FC (Nseed) RBM22High/RBM22Low
high_fidelity
-log10(p-value)
FC (Nseed) TUBA1AHigh/TUB1ALow
B
high_fidelity
-log10(p-value)
FC (Nseed) DDX3XHigh/DDX3XLow
high_fidelity
-log10(p-value)
FC (Nseed) DDX21High/DDX21Low
high_fidelity
-log10(p-value)
FC (Nseed) hnRNPA1High/hnRNPA1Low
high_fidelity
-log10(p-value)
FC (Nseed) hnRNPH2High/hnRNPH2Low
C
high_fidelity
-log10(p-value)
FC (Nseed) FUSHigh/FUSLow
high_fidelity
-log10(p-value)
FC (Nseed) hnRNPH1High/hnRNPH1Low

**Figure S5. Alternative Drosha cleavage is subjected to cellular regulation**

(A) Volcano plot of the changes in mature miRNA Number of 5' isomiRs (Nseed) between BRCA primary tumors with high/low levels (N=100 samples/each group) of control genes RBM22 and TUBA1A. (B) Volcano plot of the changes in mature miRNA Number of 5' isomiRs (Nseed) between BRCA primary tumors with high/low levels of RBPs (N=100 samples/each group) that promote increased levels of miRNA 5' isomiRs (DDX3X, DDX21, hnRNPA1 and hnRNPH2). (C) Volcano plot of the changes in mature miRNA Number of 5' isomiRs (Nseed) between BRCA primary tumors with high/low levels of RBPs (N=100 samples/each group) that promote reduced levels of miRNA 5' isomiRs (FUS and hnRNPH1).

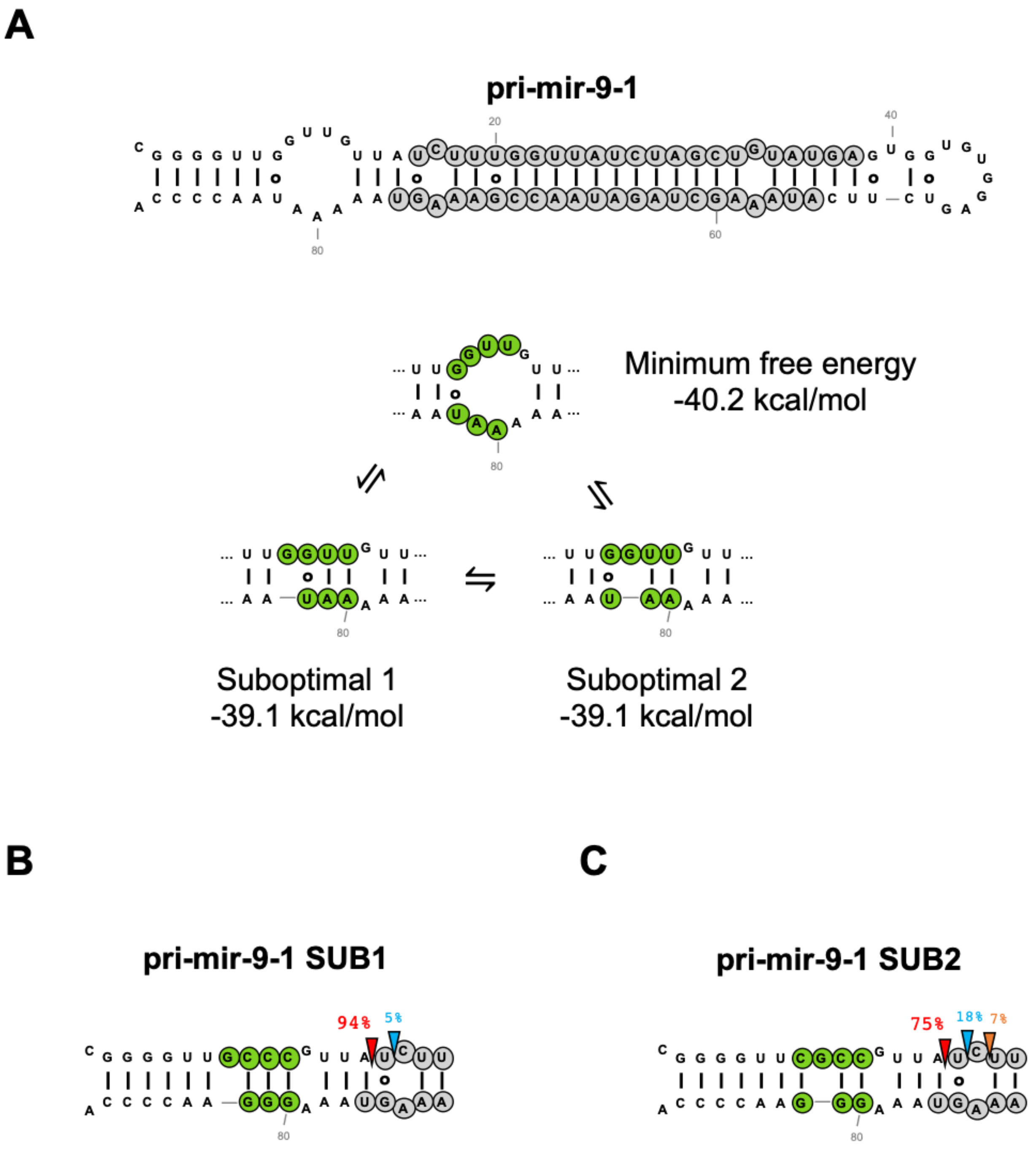
A
pri-mir-9-1
Minimum free energy
-40.2 kcal/mol
Suboptimal 1
-39.1 kcal/mol
Suboptimal 2
-39.1 kcal/mol
B
pri-mir-9-1 SUB1
94%
5%
C
pri-mir-9-1 SUB2
75%
18%
7%

**Figure S6. Suboptimal pri-miRNA conformations have distinct alternative Drosha cleavage**

(A) Scheme of conformations of wild-type pri-miR-9-1 flexible internal loop and their predicted energy. (B, C) HEK293T cells transfected with plasmids expressing pri-miR-9-1 variants that lock the local structure in the two suboptimal conformations illustrated in Supplementary Figure S6A (nucleotides changed are indicated in green). Small RNAs mapping to each pri-miRNA were used to infer the percentage of Drosha cleavage at each position.
